# Profiling the molecular signature of Satellite Glial Cells at the single cell level reveals high similarities between rodent and human

**DOI:** 10.1101/2021.04.17.440274

**Authors:** Oshri Avraham, Alexander Chamessian, Rui Feng, Lite Yang, Alexandra E. Halevi, Amy M. Moore, Robert W. Gereau, Valeria Cavalli

**Affiliations:** Department of Neuroscience, Washington University School of Medicine, St Louis 63110, Missouri, USA; Washington University Pain Center and Department of Anesthesiology, Washington University School of Medicine, St Louis 63110, Missouri, USA; Department of Neurology, Washington University School of Medicine, St Louis 63110, Missouri, USA; Neuroscience Program, Washington University School of Medicine, St Louis 63110, Missouri, USA; Department of Plastic and Reconstructive Surgery, Washington University School of Medicine, St Louis 63110, Missouri, USA; Department of Plastic and Reconstructive Surgery, The Ohio State University, Columbus Ohio, USA; Hope Center for Neurological Disorders, Washington University School of Medicine, St. Louis, Missouri 63110, USA; Center of Regenerative Medicine, Washington University School of Medicine, St. Louis, Missouri 63110, USA

**Keywords:** Satellite glial cells, DRG, pain, regeneration, human, mouse, rat, single cell RNAseq

## Abstract

Peripheral sensory neurons located in dorsal root ganglia relay sensory information from the peripheral tissue to the brain. Satellite glial cells (SGC) are unique glial cells that form an envelope completely surrounding each sensory neuron soma. This organization allows for close bi-directional communication between the neuron and it surrounding glial coat. Morphological and molecular changes in SGC have been observed in multiple pathological conditions such as inflammation, chemotherapy-induced neuropathy, viral infection and nerve injuries. There is evidence that changes in SGC contribute to chronic pain by augmenting neuronal activity in various rodent pain models. SGC also play a critical role in axon regeneration. Whether findings made in rodent model systems are relevant to human physiology have not been investigated. Here we present a detailed characterization of the transcriptional profile of SGC in mouse, rat and human at the single cell level. Our findings suggest that key features of SGC in rodent models are conserved in human. Our study provides the potential to leverage rodent SGC properties and identify potential targets in humans for the treatment of nerve injuries and alleviation of painful conditions.

## 1. INTRODUCTION

Peripheral sensory neurons located in dorsal root ganglia (DRG) relay sensory information from the peripheral tissue to the brain. Satellite glial cells (SGC) are unique glial cells in that they form an envelope that completely surround each sensory neuron [25; 57–59]. This organization allows for close bidirectional communication between SGC and their enwrapped soma. SGC actively participate in the information processing of sensory signals [28]. Morphological and molecular changes are elicited in SGC by pathological conditions such as inflammation, chemotherapy-induced neuropathic pain as well as nerve injuries [1; 9; 23; 24; 31; 80]. These studies also point to the contribution of SGC to abnormal pain conditions under injurious conditions [19; 25; 28].

Our recent analysis of SGC at the single cell level revealed that SGC share functional and molecular features with astrocytes [1; 2]. Despite great morphological differences, SGC and astrocytes share many signaling mechanisms, including potassium buffering via the inwardly rectifying potassium channel Kir4.1 and intercellular signaling via gap junctions [26]. Both cell types also undergo major changes under pathological conditions, which can have neuroprotective function but can also contribute to disease and chronic pain [26].

Most of the available information on SGC function has been obtained in rodents, which limits the potential for clinical translation. Only a small number of studies have investigated the molecular characteristics of human SGC [21]. Some evidence points to a role of SGC in human pathological conditions. In patients with Friedreich Ataxia, an autosomal recessive neurodegenerative disease, SGC proliferate, form gap junctions and abnormal multiple layers around the neurons [33; 34], likely leading to alterations in the bidirectional communication between SGC and neurons. SGC also play a role in viral infection such as herpes simplex virus or varicella zoster virus [16; 83]. In HIV-1 infection of macaques, the virus that causes AIDS, in which peripheral neuropathy and pain are common, an upregulation of GFAP in SGC was observed [44]. Hemagglutinating encephalomyelitis virus belongs to the family of coronavirus and was shown to replicate within rat sensory neurons and accumulate in lysosome-like structures within SGC, suggesting that SGC may restrict the local diffusion of the virus [40]. DRG sensory neurons and their SGC coat represent thus a potential target for multiple viral invasions in the peripheral nervous system.

A better understanding of SGC responses to mechanical, chemical and viral insults and how SGC communicate with sensory neurons will be important for future targeted therapies to treat pathological nerve conditions. To facilitate translation of findings in rodent models, a direct comparison with human tissues is thus needed. Here we present a single cell level analysis of SGC in human, mouse and rat. We find that some of the key features of SGC, including their similarities with astrocytes and the enrichment of biological pathways related to lipid metabolism and PPARα signaling are largely conserved between rodent and human. We also find notable differences in ion channels and receptors expression, which may suggest differences in SGC-neuron communication and function in painful conditions and other peripheral neuropathies. Our study highlights the potential to leverage on rodent SGC properties and unravel novel mechanisms and potential targets for treating human nerve injuries and other pathological conditions.

## 2. MATERIALS AND METHODS

### Animals and procedures

All animals were approved by the Washington University School of Medicine Institutional Animal Care and Use Committee (IACUC) under protocol A3381-01. All experiments were performed in accordance with the relevant guidelines and regulations. All experimental protocols involving rats and mice were approved by Washington University School of Medicine (protocol #20180128). Mice and rats were housed and cared for in the Washington University School of Medicine animal care facility. This facility is accredited by the Association for Assessment & Accreditation of Laboratory Animal Care (AALAC) and conforms to the PHS guidelines for Animal Care. Accreditation - 7/18/97, USDA Accreditation: Registration # 43-R-008. 8–12 week old female C57Bl/6 mice and adult male Lewis rats were used for single cell RNAseq studies.

### Single cell RNAseq in mouse and rat

L4 and L5 DRG’s from mice (2 biological independent samples, n=3 mice for each sample) and rat (2 biological independent samples, n=2 rats for each sample) were collected into cold Hank’s balanced salt solution (HBSS) with 5% Hepes, then transferred to warm Papain solution and incubated for 20 min in 37 °C. DRG’s were washed in HBSS and incubated with Collagenase for 20 min in 37 °C. Ganglia were then mechanically dissociated to a single cell suspension by triturating in culture medium (Neurobasal medium), with Glutamax, PenStrep and B-27. Cells were washed in HBSS + Hepes +0.1%BSA solution, passed through a 70-micron cell strainer. Hoechst dye was added to distinguish live cells from debris and cells were FACS sorted using MoFlo HTS with Cyclone (Beckman Coulter, Indianapolis, IN). Sorted cells were washed in HBSS + Hepes+0.1%BSA solution and manually counted using hemocytometer. Solution was adjusted to a concentration of 500cell/microliter and loaded on the 10X Chromium system. The mouse data set used in this study is the naïve data set used in our previous study [2].

Single-cell RNASeq libraries were prepared using GemCode Single-Cell 3’ Gel Bead and Library Kit (10x Genomics). A digital expression matrix was obtained using 10X’s CellRanger pipeline (Build version 3.1.0) (Washington University Genome Technology Access Center). Quantification and statistical analysis were done with Partek Flow package (Build version 9.0.20.0417). Filtering criteria: Low quality cells and potential doublets were filtered out from analysis using the following parameters: total reads per cell: 600–15000, expressed genes per cell: 500–4000, mitochondrial reads <10%. A noise reduction was applied to remove low expressing genes < = 1 count. Counts were normalized and presented in logarithmic scale in CPM (count per million) approach. We applied variance stabilizing transformation to count data using a regularized Negative Binomial regression model (Seurat::SCTransform) followed by a removal of unwanted variation caused by known nuisance and/or batch factors (Scale expression). Principal component analysis (PCA) using Louvain clustering algorithm was then undertaken followed by an unbiased clustering (Graph-based clustering) algorithm implemented in Partek. Clustering was performed using Compute biomarkers algorithm, which compute the genes that are expressed highly when comparing each cluster. Seurat3 integration was used to obtain cell type markers that are conserved across samples and clusters were assigned to a cell population by at least three established marker genes. Clusters are presented in t-SNE (t-distributed stochastic neighbor embedding) plot, using a dimensional reduction algorithm that shows groups of similar cells as clusters on a scatter plot. Differential gene expression analysis performed using three different models: Compute biomarkers following regularized Negative Binomial regression, non-parametric ANOVA and the Partek algorithm GSA that integrate multiple statistical models. Gene lists from each statistical model were intersected to remove potentially false positive genes. The intersected listed were then applied for all downstream analyses. A gene was considered differentially expressed (DE) if it has a false discovery rate (FDR) step-up (p value adjusted). p ≤ 0.05 and a Log2fold-change ≥ ± 2. The DE genes were subsequently analyzed for enrichment of GO terms and the KEGG pathways using Partek flow pathway analysis. Partek was also used to generate figures for t-SNE and scatter plot representing gene expression.

### Human tissue collection

For snRNAseq and TEM, human dorsal root ganglia (hDRG) were obtained from Anabios, Inc (San Diego, CA) (donor #1), or Mid-America Transplant (St. Louis, MO) (donors #2-5). L4-L5 DRG were extracted from tissue/organ donors less than 2 hours after aortic cross clamp and was immediately snap-frozen and stored at −80C until use. Details regarding the donors for snRNAseq are included below:

| Donor # | Age | Sex | Race | DRG level | BMI | COD |
| --- | --- | --- | --- | --- | --- | --- |
| 1 | 44 | Female | Asian | L4 |  | CVA |
| 2 | 56 | Female | Black | L5 | 23.13 | Anoxia/OD |
| 3 | 23 | Male | White | L5 | 21.37 | Head trauma/MVA |
| 4 | 45 | Male | White | L4,5 | 28.8 | Anoxia |
| 5 | 51 | Female | White | L5 | 33.49 | Anoxia |

For immunohistochemistry studies, human DRG were obtained from organ donors with full legal consent for use of tissue in research, and in compliance with procedures approved by Mid-America Transplant (St. Louis, MO). The Human Research Protection Office at Washington University in St. Louis provided an Institutional Review Board waiver. Details regarding the donors for immunohistochemistry are included below:

| Donor # | Age | Sex | Race | DRG level | BMI | COD |
| --- | --- | --- | --- | --- | --- | --- |
| 6 | 46 | Female | White | L1,2 | 20.91 | Anoxia |
| 7 | 44 | Female | Middle Eastern | L1,4 | 22.2 | CVA/SAH |
| 8 | 53 | Female | White | L4,5 | 25.48 | CVA/Stroke |
| 9 | 54 | Female | White | L1-5 | 25.9 | CVA/Stroke |

### Single nucleus RNAseq from human sample

To make the tissue suitable for nuclei isolation, the entire DRG was processed into smaller pieces by cryopulverization using the CryoPrep (Covaris; CP02). Nuclei were isolated according to Martelotto with some modifications [45]. We elected to apportion the cryopulverized tissue and process the portions in parallel using two different buffers in order to evaluate potential effects on nuclei representation. One homogenization buffer was EZ Nuclei Lysis Buffer (Sigma; NUC101-1KT) with 0.5% RNasin Plus (Promega; N2615), 0.5% SUPERase-In (ThermoFisher; AM2696) and 1mM DTT [22]. The other homogenization buffer was CST buffer (NaCl2 146 mM, Tris HCl pH 7,5 10 mM, CaCl2 1mM, MgCl2 21 mM, 0.49% CHAPS (Millipore-Sigma), 0.01% BSA, 0.5% SUPERasin-in, 0.5% RNasin Plus), as described in Slyper et al. [68]. Using the EZ buffer, the samples were homogenized on ice using 18 strokes of Pestle A followed by 18 strokes of Pestle B. The homogenate was filtered through a 50 *m.u*m filter (Sysmex; 04-004-2327) into a 2 mL microcentrifuge tube (Eppendorf; 022431048). An additional 0.5 mL of homogenization buffer was used to wash the Dounce homogenizer and filter. The sample was then placed on ice while the remaining samples were processed. The sample was centrifuged at 500g at 4°C for 5 mins to obtain a crude pellet containing spinal nuclei. The supernatant was removed and discarded, being careful to not disturb the pellet. The pellet was resuspended in 1.5 mL of Homogenization Buffer and allowed to sit on ice for 5 mins. at 500g, 4°C for 5 mins. This wash step was repeated twice more for a total of 3 washes. The final pellet was resuspended in 0.5 mL of NRB containing 6 *μ*M 4’,6-diamidino-2-phenylindole (DAPI, ThermoFisher; D1306). The suspension was filtered through a 20 *μ*m filter (Sysmex; 04-004-2325) into a polypropylene tube and kept on ice. Using the CST buffer, the samples were homogenized on ice using 18 strokes of Pestle A followed by 18 strokes of Pestle B in 1 mL of CST buffer. The homogenate was filtered through a 50 um filter into a 15 mL conical. An additional 1 mL was used to wash the filter and then 3 mL of CST was added, bringing the total volume to 5 mL. The suspension was spun down at 500g for 5 mins at 4°C. The supernatant was removed and the pellet was resuspended in 0.5 mL of CST containing 6 *μ*M of DAPI. The suspension was filtered through a 20 *μ*m filter into a polypropylene tube and kept on ice. Fluorescence Activated Nuclear Sorting (FANS) was performed to purify nuclei from debris on a FACSAria II (BD). Gates were set to isolate DAPI+ singlet nuclei based on forward scatter and side scatter as well as fluorescence intensity. The instrument was set to 45 pounds per square inch (psi) of pressure and a 85 *μm* nozzle was used, with sterile PBS sheath fluid. Nuclei were sorted into a 1.5 ml microcentrifuge tube containing 15 *μl* of NRB at 4°C. For each sample, 18,000 events were sorted into the collection tube. The sorted nuclei and NRB total volume was approximately 45 ul, allowing for the entire loading of the suspension into the Chromium Single Cell 3’ v3 solution (10x Genomics) without any further manipulation. 10x libraries were processed according to the manufacturer’s instructions. Completed libraries were run on the Novaseq 6000 (Illumina). A digital expression matrix was obtained using 10X’s CellRanger pipeline as above.

### Tissue preparation and immunohistochemistry

Mice were perfused with PBS buffer, followed by 4% paraformaldehyde. After isolation of mouse DRG, the tissue was post-fixed using 4% paraformaldehyde for 1 h at room temperature. Tissue was then washed in PBS and cryoprotected using 30% sucrose solution at 4°C overnight. Next, the tissue was embedded in O.C.T., frozen, and mounted for cryosectioning. Mouse frozen sections were cut at 12 μm for subsequent staining. Freshly dissected human DRG were sectioned to 40 μm, fixed and stored as free floating in cryoprotectant. Mouse DRG sections mounted on slides and human floating DRG sections were washed 3x in PBS and then blocked in solution containing 10% donkey serum in 0.1% Triton-PBS for 1 h. Next, sections were incubated overnight in blocking solution containing primary antibody. The next day, sections were washed 3x with PBS and then incubated in blocking solution containing a secondary antibody for 1 h at room temperature. Finally, sections were washed 3x with PBS and mounted using ProLong Gold antifade (Thermo Fisher Scientific). Images were acquired at 10x or 20x using a Nikon TE2000E inverted microscope. Antibodies were as follow: TUJ1 (Tubb3/βIII tubulin) antibody (BioLegend catalog #802001, RRID:AB_291637), FABP7 (Thermo Fisher Scientific Cat #PA5-24949, RRID:AB_2542449), FASN (Abcam, Catalog #ab128870). Stained sections with only secondary antibody were used as controls. Quantification of SGC markers in human and mouse DRG sections were done in ImageJ where the % of neurons surrounded by at least one SGC expressing the indicated markers out of total number of neurons in each section was quantified. n=4 biological independent replicates. Unpaired t-test. Data are presented as mean values ±SD.

### Transmission electron microscopy of mice and human DRG

Mice were perfused with 2.5% glutaraldehyde with 4% paraformaldehyde in 0.1M cacodylate buffer and DRG were drop fixed to the same fixation buffer for a post fix. A secondary fix was done with 1% osmium tetroxide. Freshly collected human DRG samples were drop fixed in 2.5% glutaraldehyde + 2% paraformaldehyde in 0.15M (final concentration) cacodylate buffer pH 7.4 with 2mM calcium chloride overnight at 4C. Samples were then vibratomed in the same buffer used for fixation and sections collected into buffer in well plates. For Transmission electron microscopy (TEM), tissue was dehydrated with ethanol and embedded with Spurr’s resin. Thin sections (70 nm) were mounted on mesh grids and stained with 8% uranyl acetate followed by Sato’s lead stain. Sections were imaged on a Jeol (JEM-1400) electron microscope and acquired with an AMT V601 digital camera. (Washington University Center for Cellular Imaging).

## 3. RESULTS

### Profiling SGC from human, mouse and rat at the single cell level

To define the similarities and differences between SGC across different species, we performed snRNA seq of L4,L5 human DRG and scRNA-seq of L4,L5 mouse and rat DRG using the Chromium Single Cell Gene Expression Solution (10x Genomics) (Fig. 1A). We chose to perform scRNAseq in rodents because we previously showed that this method efficiently captures SGC [1; 2]. We opted for snRNAseq in human DRG because the tissue was frozen and the large size of cells in human may limit their capture rate in the 10x platform. The number of sequenced human nuclei from five donors (donor information is in the methods section) was 19,865, with an average of 129,520 mean reads per cell, 1,480 mean genes per cell and a total of average 25,643 genes detected. The number of sequenced mouse cells from two independent biologically replicates (pooled DRG’s from 3 mice for each replicate) was 6,343 with an average of 65,378 mean reads per cell, 1,510 mean genes per cell and a total of 18,130 genes detected. The number of sequenced rat cells from two biologically independent replicates (pooled DRG’s from 2 rats for each replicate) was 15,892 with an average of 41,594 mean reads per cell, 2,132 mean genes per cell and a total of 17,137 genes detected. Low quality cells and doublets were filtered out from downstream analysis (see filtering criteria in the methods). Cells from different donors clustered together by cell type, with the exception of donor #3, in which SGC clustered separately from the other 4 donors (Fig 1B). Batches from mouse and rat demonstrated high similarities in cell clustering (Fig. 1B).

**Figure 1:**
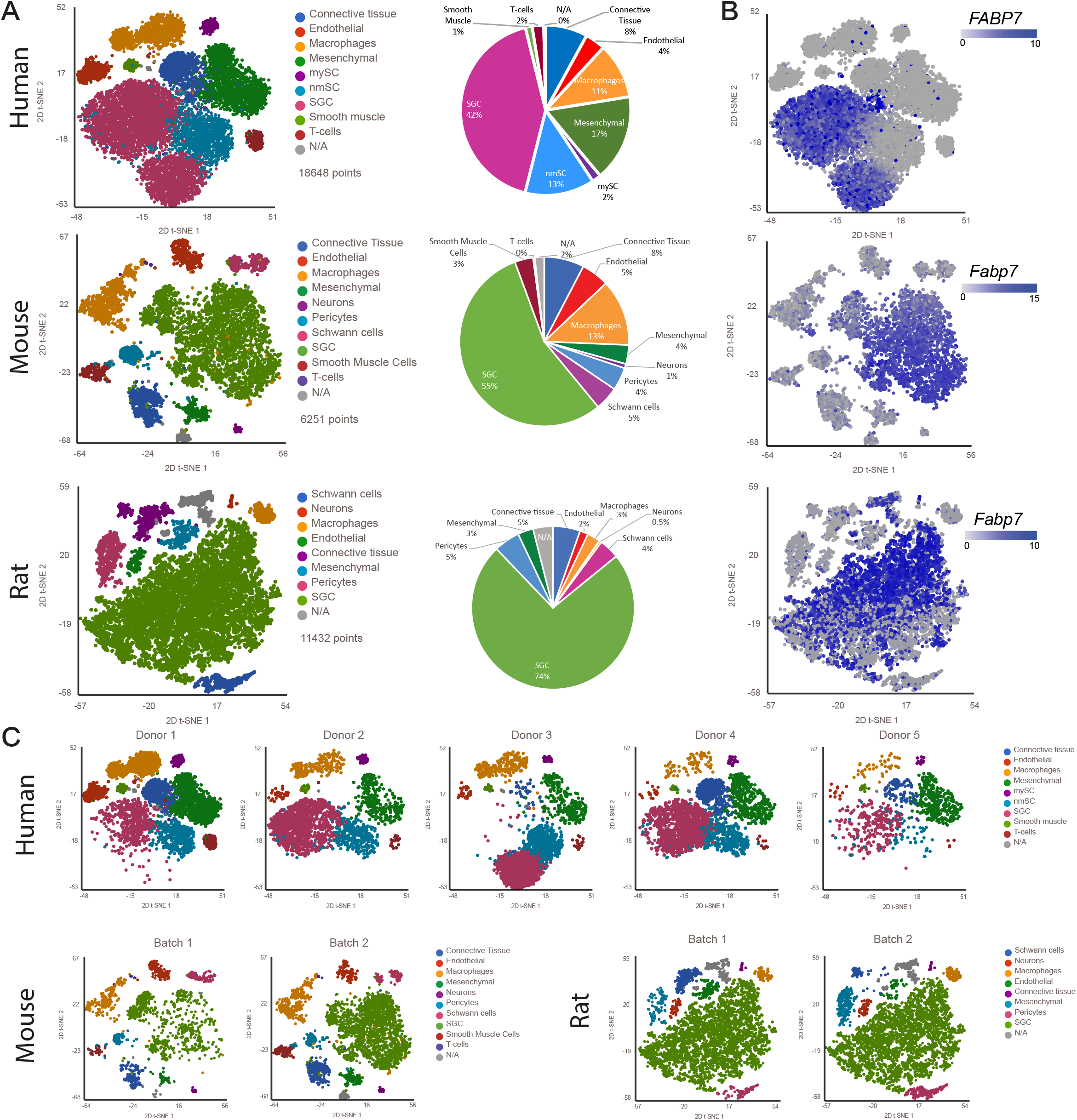
Profiling DRG cells from mouse, rat and human. (A) t-SNE plot of human (18,648 cells), mouse (6,251 cells) and rat (11,432 cells) DRG cells colored by cell populations with fraction of distribution (B) t-SNE plots separated by batch/donors of human (donor1 6,444 cells, donor2 3,498 cells, donor3 3,957 cells, donor4 3,871 cells and donor5 878 cells), mouse (batch1 1,929 cells batch2 4,322 cells) and rat (batch1 6,074 cells batch2 5,358 cells) (C) t-SNE overlay for expression of the SGC marker gene *Fabp7* in human, mouse and rat

To identify cluster-specific genes, we calculated the expression difference of each gene between that cluster and the average in the rest of the clusters (ANOVA fold change threshold >1.5). Examination of the cluster-specific marker genes revealed major cellular subtypes including neurons, SGC, endothelial cells, Schwann cells, pericytes, smooth muscle cells, macrophages, mesenchymal cells and connective tissue cells (Fig. 1A, table 1) [1]. Human specific marker genes were used to classify cell populations; macrophages (*CD163,MRC1*), Mesenchymal cells (*APOD,PDGFRA*), endothelial cells (*FLT1,PECAM,CLDC5*), connective tissue/mesenchymal (*COL1A1,DCN*), Myelinating Schwann cells (*PRX,MAG,PMP22*), SGC (FABP7,CDH19), Myelinating/non-Myelinating Schwann cells and SGC (*S100B*), T-cells (*Cd2,Cd3g,Cd28*) and Smooth muscle cells (*MYOCD,ACTA2,DES*) (Supplementary Fig. 1A and Table 1). Rodents had slightly different cell types and markers; macrophages (*Cd68,Aif1*), pericytes (*Kcnj8,Pdgfrb*), Neurons (*Tubb3,Gal,Tac,Prph*), SGC (*Cdh19,Fabp7,Kcnj10*), Schwann cells (*Prx,Mag,Pmp2*), Mesenchymal cells (*Apod,Pdgfra*), Smooth muscle cells (*Myocd,Acta2,Des*), endothelial cells (*Flt1, Pecam, Cldn5*), connective tissue (*Col1a1,Dcn*) (Supplementary Fig. 1B,C and Table 1). The cell clusters obtained from the mouse data set matched our previous dataset [1] (Supplementary Fig. 1D).

**Table 1.**
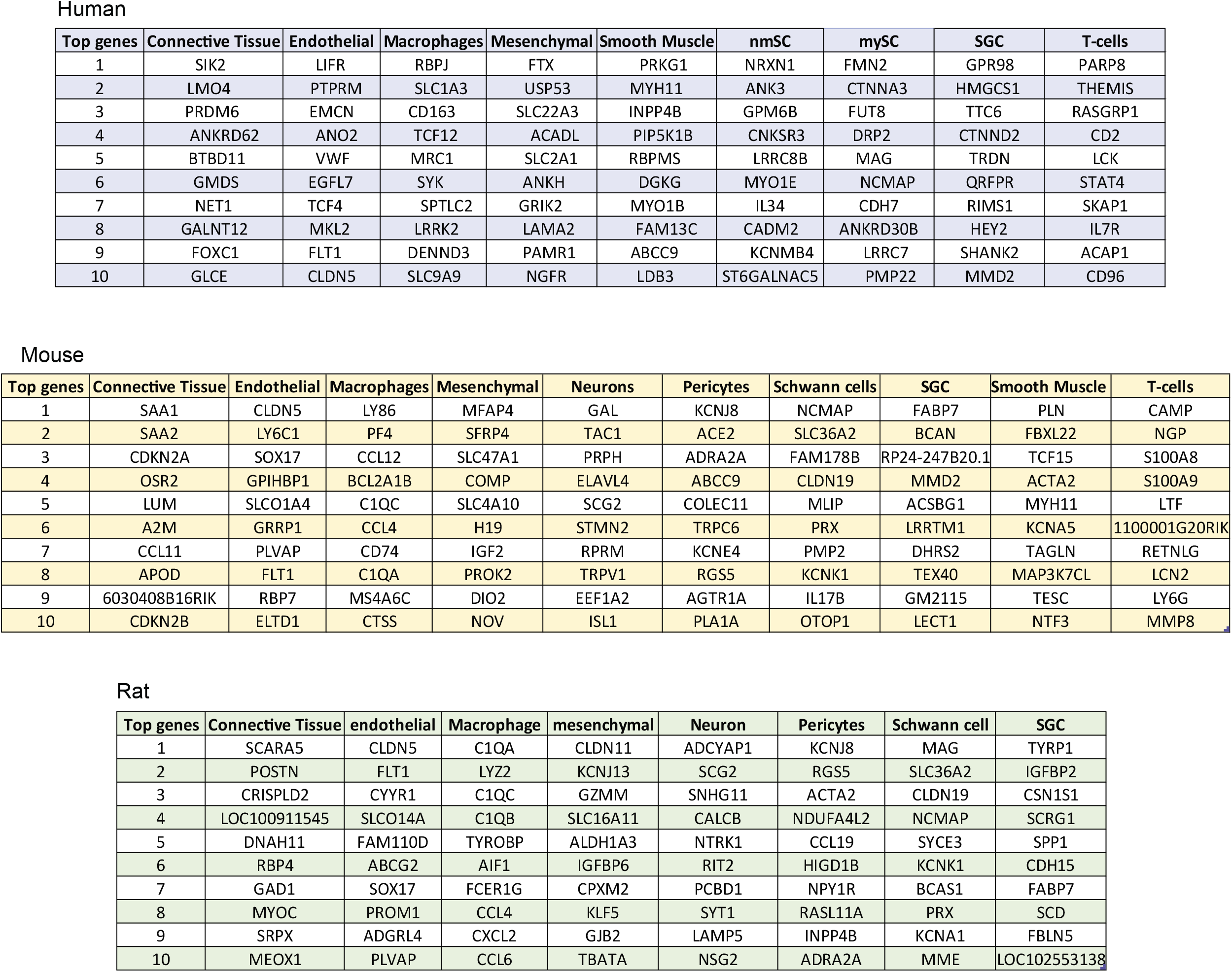
Top 10 differentially expressed genes in each cell population compared to all other cell types in the DRG (fold change threshold >1.5)

We previously showed that although the actual percentage of neuronal cells in the mouse DRG is about 12%, the number of neurons detected in our scRNAseq analysis was only about 1% [1; 2], which might be a result of the dissociation protocol that is biased towards non-neuronal cells. Another possibility is neuronal damage during the tissue dissociation process or the fact that sensory neurons are relatively large cells and are less amenable for single cell studies. In the rat samples, only 0.5% of cells were neurons and no neuronal cells were detected in the human sample (Fig. 1A). Nevertheless, our protocol achieved recovery of SGC from all species with 42% in human, 55% in mouse and 74% in rat (Fig. 1A), allowing us to compare the molecular profile of SGC across species.

We recently described that *Fabp7* (Fatty acid binding protein 7) is a specific marker gene for SGC and that the FABP7 protein is highly enriched in mouse SGC compared to other cells in the DRG [1]. t-SNE plots overlaid for *Fabp7* demonstrated that *Fabp7* is also enriched in human and rat SGC (Fig. 1C). To validate *Fabp7* expression at the protein level, we performed immunostaining of DRG sections from mouse and human, which revealed specific FABP7 labelling of SGC surrounding sensory neurons in both species (Fig. 2A,B).

**Figure 2:**
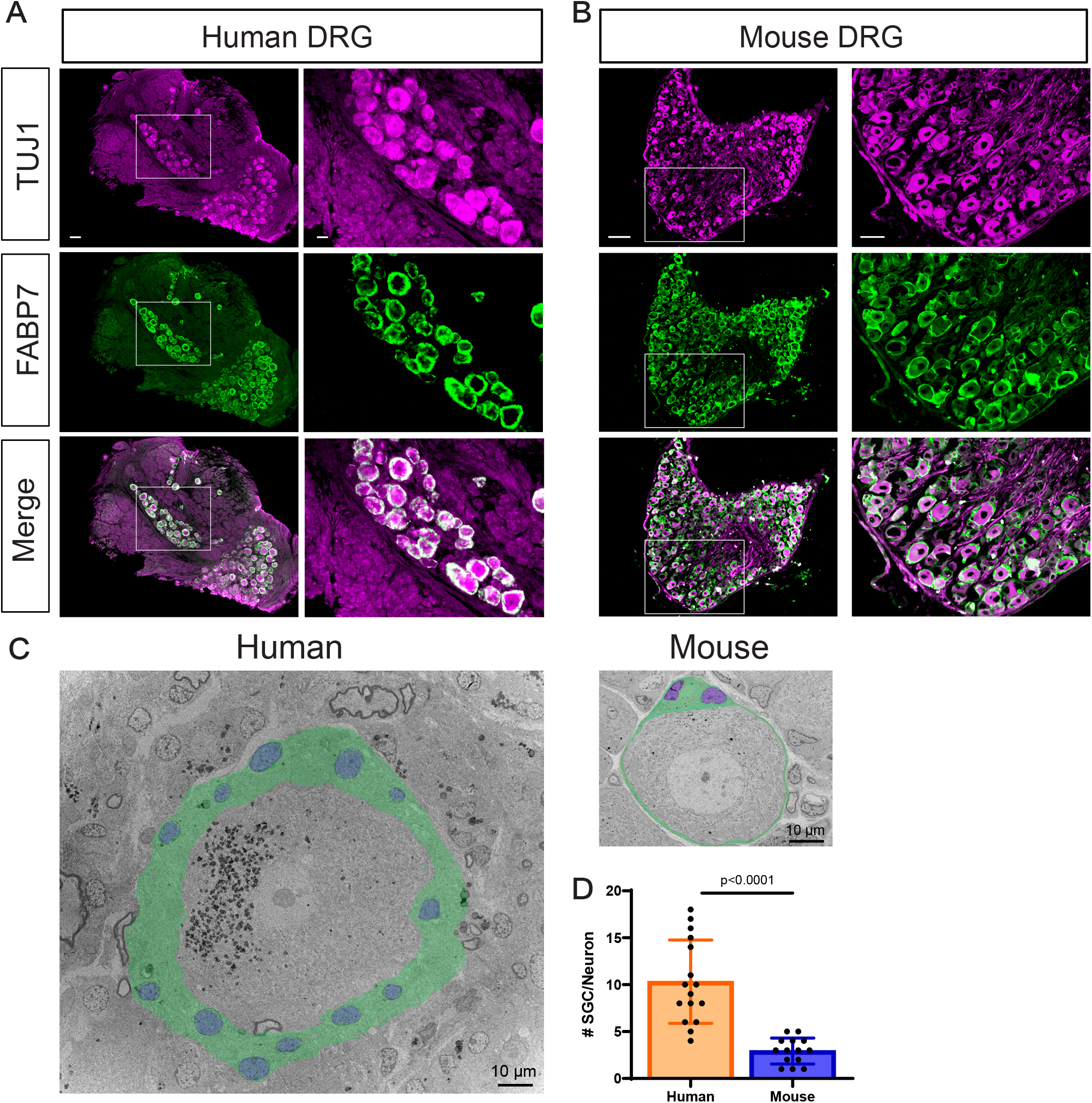
FABP7 is an SGC marker in mouse and human. (A and B) Representative images of immunofluorescence staining of human (a) and mouse (b) DRG sections labelled for TUJ1 (magenta) and FABP7 (green) Scale bar: 200μm (left panel) 50μm (right panel). n=4 biological independent replicates for both a and b. (C) Representative TEM images of DRG sections from human and mouse showing neuronal cell bodies and its enveloping SGC (SGC cytoplasm is pseudocolored in green, SCG nuclei are pseudocolored in blue). (D) Quantification of the number of SGC surrounding a single neuron. n=4 biologically independent animals (mouse), n=2 individual donors (human).

In adult animals, SGC tightly enwrap the soma of each sensory neuron [56; 57; 59]. The gap between SGC and the neuronal surface is only about 20 nm, which is similar to that of the synaptic cleft. This close association between the two cell types is essential for efficient mutual neuron–SGC interactions [25]. The detailed morphology of the human neuron-SGC unit has only been examined at the light microscopy level in human [21]. To further compare the SGC organization surrounding sensory neurons across species, we performed Transmission Electron Microscopy (TEM) of human and mouse DRG sections (Fig. 2C), which demonstrated the tight contact between SGCs and neurons in both mouse and human and the increased human sensory neuron soma size, which can be up to 5 times larger than mouse sensory neurons (Fig 2C) [21]. Quantification of the number of SGC surrounding sensory neurons revealed that human sensory neurons are surrounded by significantly more SGC compared to mouse neurons (Fig. 2D), consistent with the observations that the number of SGC surrounding sensory neurons increases with increasing soma size in mammals [36; 56; 59].

### Expression of SGC specific marker genes in rodents and human

We next examined the expression of known SGC marker genes in rodent and human. Cells in the clusters identified as SGC were pooled together. We identified 7,880 SGC in human, 3,460 SGC in mouse and 8,428 SGC in rat (Fig. 3A-C). t-SNE plots overlaid with *Fabp7* demonstrated that in all species, *Fabp7* is expressed at high levels in a majority of SGC (mouse 92%, rat 88% and human 96% (Fig.3A-D). Cadherin19 (*Cdh19*) has been described as a unique SGC marker in rat Schwann cell precursors [71] and in adult rat SGC [20]. We found that *Cdh19* was expressed in most human SGC (98%), whereas only half of SGC in rodents expressed this gene (56% in mouse and 48% in rat (Fig. 3A-D). Glutamine synthetase (*GS/Glul*) has been suggested as an SGC specific marker in rat [46] and mouse DRG [30; 31]. Our previous finding indicated a nonspecific expression of *Glul* in almost all cells in the DRG at the transcript level [1]. Our current analysis suggests differences in *Glul* expression between species with more than half of SGC expressing *Glul* in rodents (~60%), and only around 10% in human (Fig. 3A-D).

**Figure 3:**
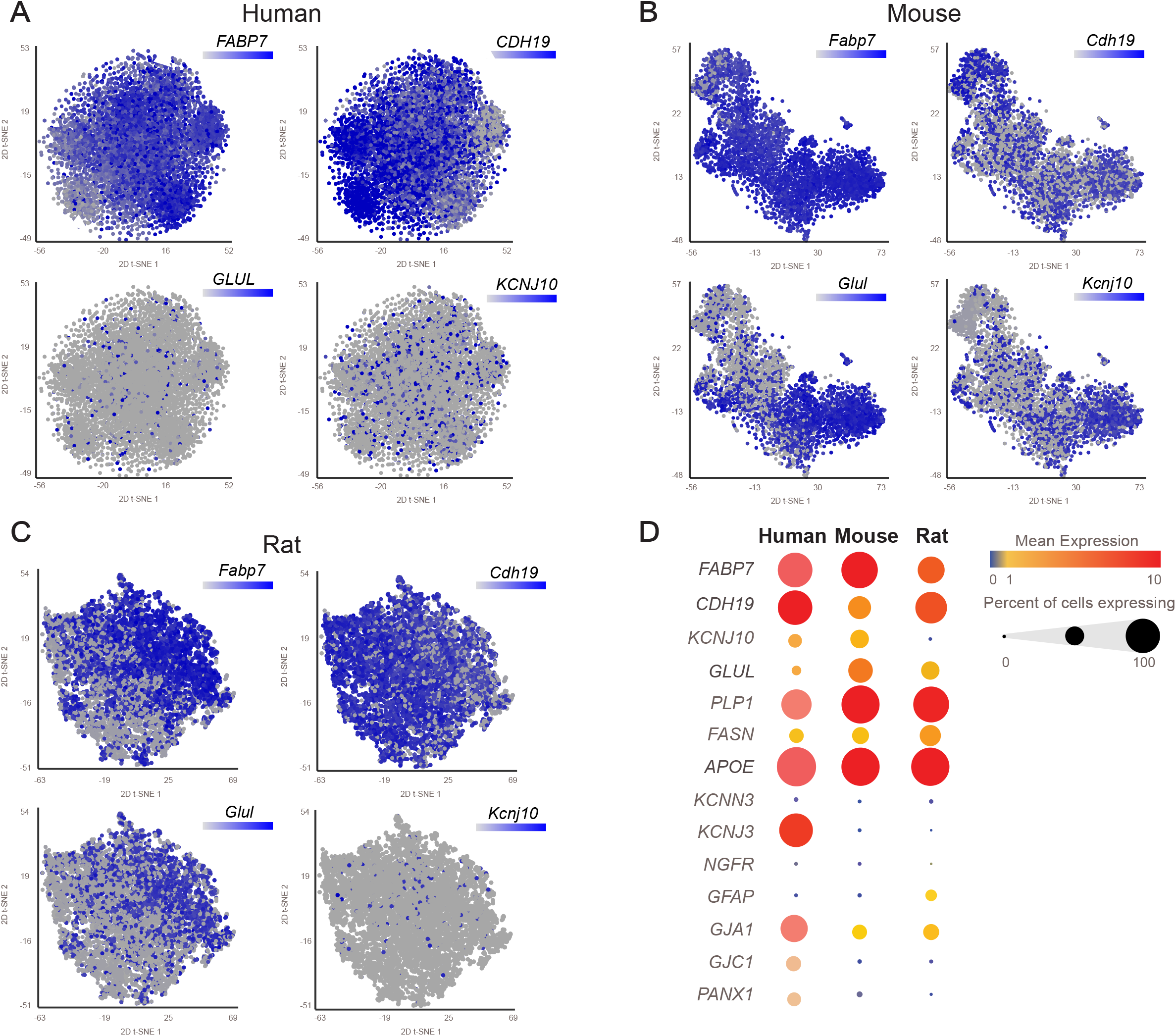
SGC marker genes in mouse, rat and human. (A) t-SNE overlay of pooled SGC for expression of SGC marker genes *FABP7, CADH19, GLUL* and *KCNJ10* in human (7,880 cells) (B) t-SNE overlay for expression of SGC marker genes *Fabp7, Cadh19, Glul* and *Kcnj10* in mouse (3,460 cells) (C) t-SNE overlay for expression of SGC marker genes *Fabp7, Cadh19, Glul* and *Kcnj10* in rat (8,428 cells) (D) Bubble plot showing the normalized count expression of the indicated SGC marker genes in each species. Dot size represents the percentage of cells that express the corresponding genes and the color represents the mean expression level.

One of the characteristics of SGC, which is similar to astrocytes, is the ability to control the microenvironment via expression of transporters and ion channels [26]. The potassium channel Kir4.1/*Kcnj10* is a known marker of SGC. Kir4.1/*Kcnj10* is expressed in rat SGC and influence the level of neuronal excitability, which has been associated with neuropathic pain conditions [79]. Our analysis demonstrates that *Kcnj10* is expressed in almost half of mouse SGC (42%), whereas it is expressed in fewer SGC in rat (6%) and human (17%) (Fig.3A-D). Interestingly, we found that the potassium channel Kir3.1/*Kcnj3* is widely expressed in human (82%) but not in rodents SGC (Fig. 3D). The diversity in potassium channel expression in SGC might suggest a special role for this channel in defining the physiological characteristics of SGC across species. Another potassium channel that has been shown to be expressed specifically in rat SGC is SK3/*Kcnn3* [79]. Our analysis suggests that only a small subset of rat SGC (12%) and human SGC (6%) express *Kcnn3* (Fig.3D), whereas mouse SGC do not express *Kcnn3* (Fig.3D).

Another main property of SGC that is shared with astrocytes is functional coupling by gap junctions, with SGC surrounding the same neuron connected by gap junctions [25; 29]. Gap junction protein alpha 1 (*CX43/Gja1*) is the most abundant connexin (Cx) and was shown to modulate pain responses [69]. We found that *Gja1* is expressed in a majority of human SCG (70%) but less prevalent in mouse (30%) and rat (40%) (Fig. 3D). Other known gap junctions proteins expressed in SGC include Cx32, followed by Cx30.2, Cx37, Cx26, Cx30, Cx45 and Cx36 [25]. Although these gap junction genes were reported to be expressed in SGC, we found that less than 5% of SGC expressed them across all species, except for *Cx30.2/Gjd3* which was expressed in 35% of mouse SGC and *Cx45/Gjc1* that is expressed in 30% of human SGC (Fig. 3D).

Membrane channels related to gap junctions are Pannexins (Panx), which do not form cell-to-cell channels but are highly permeable to ATP [28]. Pannexin1(*Panx1*) was reported to be expressed in sensory ganglia where it is increased in pain models [69] and there is evidence that *Panx1* mediated ATP release is implicated in nociception [25]. Our analysis demonstrated moderate expression of *Panx1* in human SGC (30%) with lower expression in mouse and rat (Fig.3D). These observations indicate variability in Gap junction and pannexin gene expression between rodent and human, which may suggest functional differences in SGC communication and function in nociception. SGC also express glial fibrillary acidic protein (GFAP) and similarly to astrocytes, GFAP expression is increased under pathological conditions, which can have a protective function [82]. In mouse, *Gfap* is one of the top upregulated genes in SGC upon nerve injury [1; 2; 10; 17], but it is not expressed in all SGC [2; 47]. In uninjured SGC, the distribution of *Gfap* was relatively low, with ~20% expression in rat, 1.5% in mouse and undetectable levels in human DRG (Fig. 3D).

While SGC do not typically myelinate neuronal soma, except in the spiral ganglion [62], some myelin associated genes such as *Mpz*, *Mbp*, and *Plp1* are highly expressed in SGC [1]. Proteolipid protein (*Plp1*) is the major myelin protein in the central and peripheral nervous system. *Plp1* is expressed in all rodents SGC and in more than half of human SGC (Fig.3D). Another gene that is expressed in all rodent and human SGC is apolipoprotein E (*ApoE*) (Fig.3D). APOE is a multifunctional protein, mainly involved in lipid synthesis and transport. High levels of APOE production occurs in the brain, where it is primarily synthesized by astrocytes [18]. We recently found that one of the genes enriched in mouse SGC is fatty acid synthase (*Fasn*) [1], which controls the committed step in endogenous fatty acid synthesis [14]. Examination of *Fasn* transcript expression revealed high distribution in rat SGC (70%) and lower in mouse (40%) and human (33%) SGC (Fig. 3D).

To validate the expression of unique SGC marker genes across species at the protein level, immunostaining for selected markers was performed in human and mouse DRG sections. Immunostaining for FASN revealed its specific expression in SGC surrounding neurons (stained for TUJ1) in both human and mouse DRG tissue (Fig. 4A). These results suggest that the expression of genes related to lipid metabolism and transport in SGC, such as *Fasn* are conserved between rodents and human. *Gfap* was detected only at low levels in our scRNAseq analysis and immunostaining with GFAP further demonstrated expression only in very few SGC in both mouse and human (Fig. 4B). The SGC marker *Glul* was highly expressed in mouse SGC and less in human at the RNA level (Fig. 3D). Staining for GLUL also revealed higher expression around most mouse SGC, with lower detection in human SGC (Fig.4C). Another striking difference between human and mouse SGC is the expression of the potassium channel *Kcnj3* (Kir3.1), which is highly expressed in human SGC but not in rodent (Fig 3D). Immunostaining for *Kcnj3* (Kir3.1) further confirmed expression in human SGC, with almost undetectable expression in mouse SGC (Fig. 4D). Quantification of the percent of neurons associated with SGC expressing FASN, GFAP, GLUL or KCNJ3 (Fig. 4E) confirms the important similarities and differences in ion channels and gap junction genes between human and mouse SGC that might impact their function in controlling neuronal activity and nociceptive thresholds.

**Figure 4:**
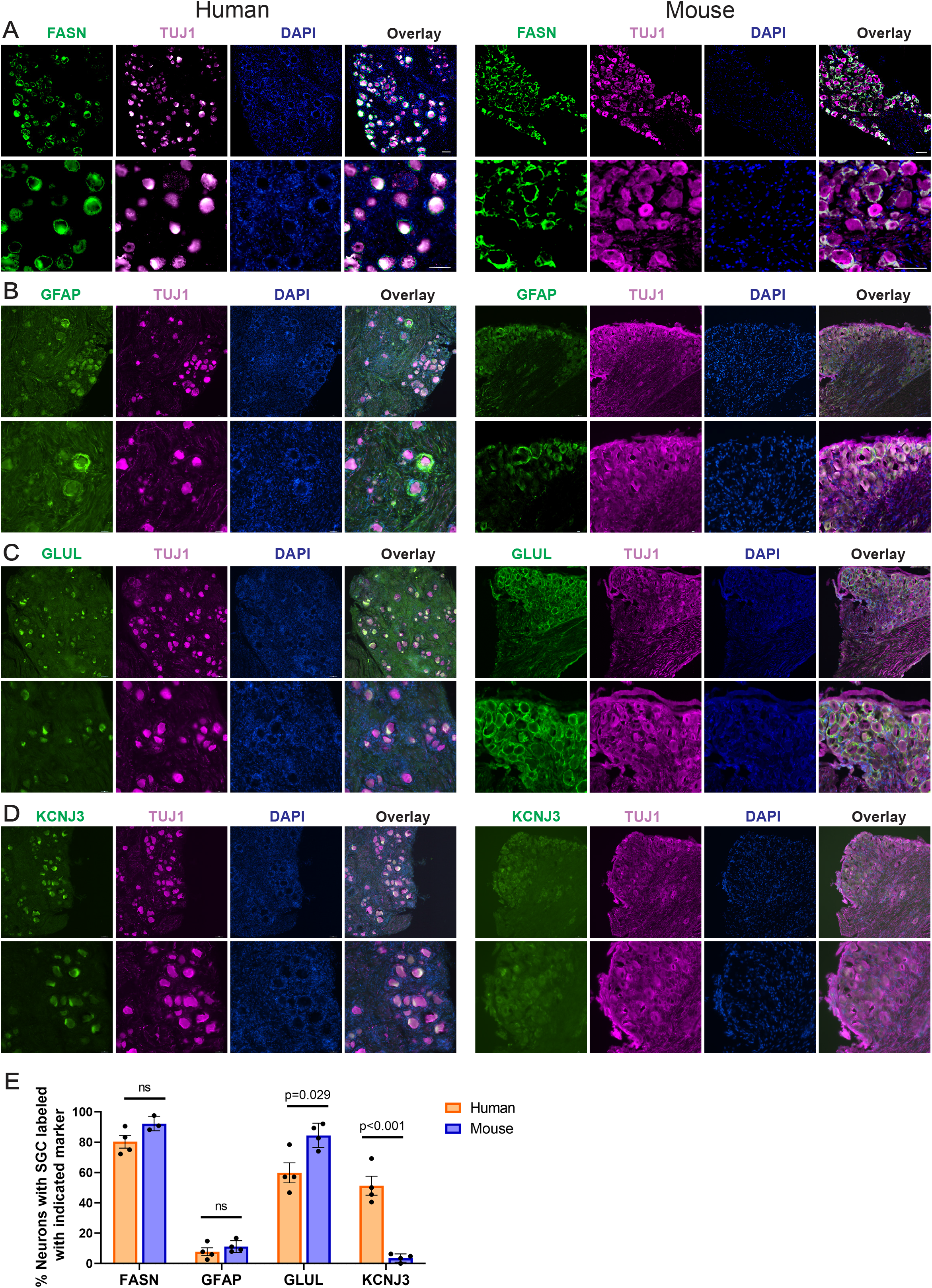
Immunostaining of SGC in human and mouse DRG sections. (A-D) Representative images of immunofluorescence staining of human and mouse DRG sections labelled with TUJ1 (magenta), DAPI (Blue) and FASN (green) (A), GFAP (B), GLUL (C), and KCNJ3 (D). Scale bar: 100μm. n=4 biological independent replicates (E) Quantification of SGC markers in human and mouse DRG sections. The % of neurons surrounded by at least one SGC expressing the indicated markers out of total number of neurons in each section was quantified. FASN; 80.280±8.325 in human and 92.235±4.802 in mouse, GFAP; 7.8±5.13 in human, 11.191±3.884 in mouse, GLUL; 59.833±13.314 in human and 84.518±8.021 in mouse, KCNJ3; 51.306±12.558 in human and 3.615±2.668 in mouse. n=4 biological independent replicates. Unpaired t-test. Data are presented as mean values ±SD.

### SGC in rodent and human share functional properties

To further examine the biological properties of SGC across species, we calculated the top differentially expressed genes in SGC in human (2,070 genes), mouse (1,622 genes) and rat (993 genes) (fold-change >1.5, significant differences across groups, and p < 0.05 compared to average gene expression in all other populations in the DRG in the same species). This analysis might be influenced by the fact that the representation of cell population differs in each data set (Fig 1A). Nonetheless this analysis allowed us to compare the three gene sets, which revealed 193 genes shared between SGC in human and rodents (Fig. 5A, Supplementary Table 1). The common genes included *Fabp7, ApoE, Fasn, Kcnj10*, and *Gja1*, suggesting conserved roles of SGC related to lipid metabolism (Fabp7, ApoE and Fasn), physiological properties (*Kcnj10*) and cell-cell communication (*Gja1*). Many of the shared genes were also expressed in astrocytes (Supplementary Fig. 2A), consistent with our previous findings that 10% of top enriched genes in mouse SGC were shared with brain astrocytes [1]. We next examined if human SGC also share unique expressed genes with human mature and fetal astrocytes [84]. We found that human SGC shared 152 genes with human mature astrocytes, 107 genes with human fetal astrocytes and 260 genes were shared with both mature and fetal astrocytes (Supplementary Fig. 2A, Supplementary Table 2). Despite the diversity in morphology and signaling mechanisms between astrocytes and SGC, these results support that important parallels between these two cell types are conserved between mice and human.

**Figure 5:**
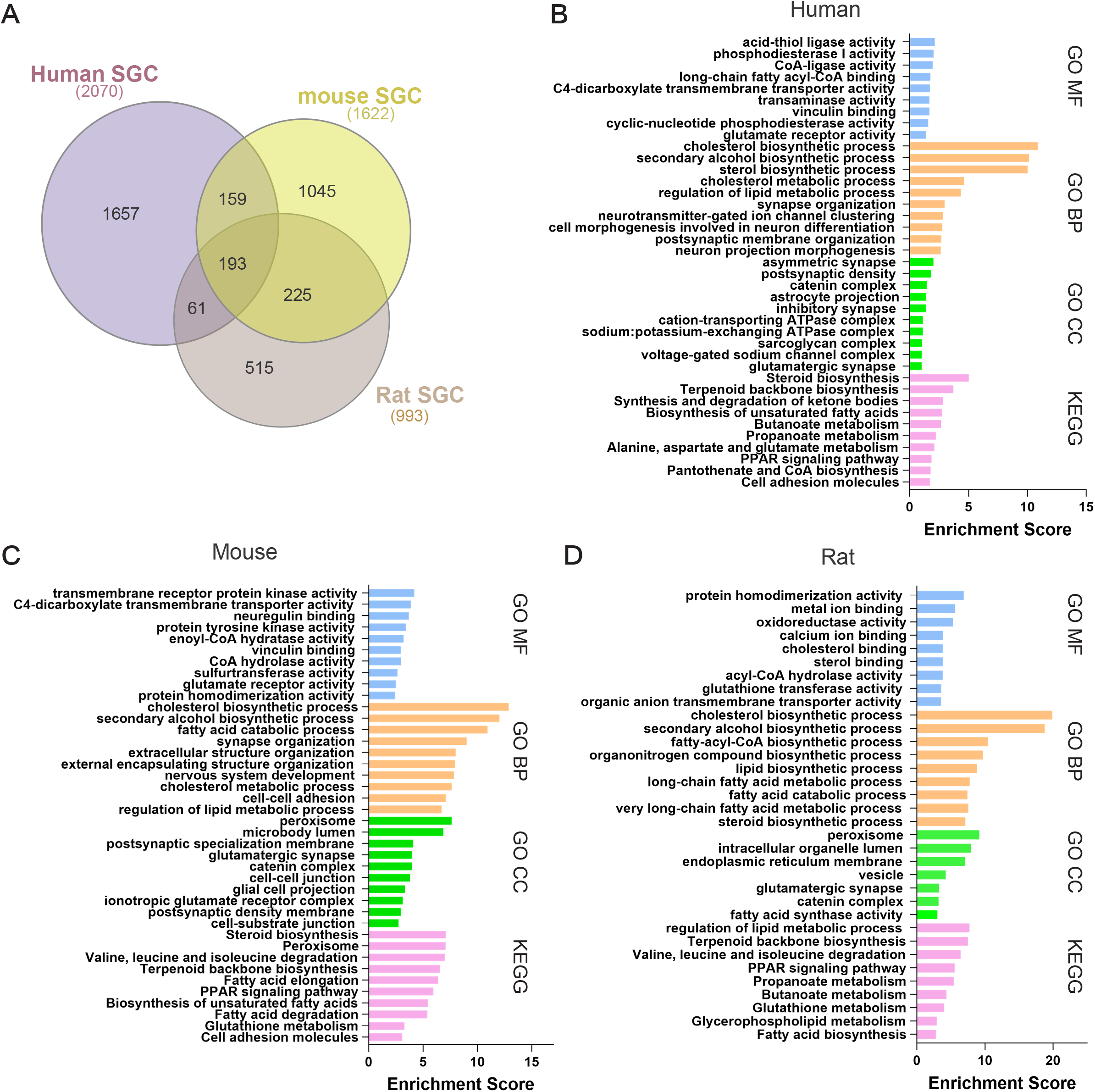
Enrichment of genes and pathways in human, mouse and rat SGC. (A) Venn diagram for enriched genes in SGC compared to other cells in the DRG from mouse (1,622 genes), rat (993 genes) and human (2,070 genes). (B) Pathway analysis (KEGG/GO) of enriched genes express in human SGC. (C) Pathway analysis (KEGG/GO) of enriched genes express in mouse SGC. (D) Pathway analysis (KEGG/GO) of enriched genes express in rat SGC.

We next compared human SGC to Schwann cells, the other main types of glial cells in the peripheral nervous system [81]. Schwann cells are divided to two types; myelinating (mySC) and non-myelinating (nmSC). We identified both types of Schwann cells in our human scRNAseq data set and compared their similarity to SGC. We found that human SGC share 453 genes with nmSC, 350 genes with mySC and 260 genes with both mySC and nmSC (Supplementary Fig. 2B, Supplementary Table 3). Comparison of Schwann cell types between mouse [81] and human showed more similarity between mouse mySC and human nmSC (Supplementary Fig. 2C, Supplementary Table 4). Our recent work also identified that mouse SGC do not represent a uniform cell population and at least four subtypes exist [2]. One of the sublcuster (mSGC3) we defined showed the highest similarity to astrocytes and another subcluster (mSGC 4) resembled the most to mySC [2]. We found that human SGC expressed a unique gene set that showed the highest similarity to the mSGC3 (Supplementary Fig. 2D, Supplementary Table 5). These results further support the high similarity at the gene expression level between SGC and astrocytes and extend this similarity to human tissue.

We next analyzed the enriched biological pathways using KEGG 2021 (Kyoto Encyclopedia of Genes and Genomes) and Gene Ontology (GO). We found that human and rodents SGC show enriched molecular functions (GO MF) related to enzymatic activity and ion channel and transport activity (Fig. 5B-D). This further supports the important role SGC may play in human DRG to control neuronal excitability and pain thresholds. Both rodents and human SGC shared some cellular components (GO CC), including involvement in plasma membrane and synaptic activity (Fig. 5B-D). The biological process enriched pathways (GO BP) in human and rodents were particularly similar, with enrichment for mainly metabolic pathways of lipids and cholesterol, as well as processes related to nervous system development (Fig. 5B-D). We previously revealed that fatty acid synthesis and PPARα signaling pathway were enriched in mouse SGC and those pathways were also upregulated after peripheral nerve injury in SGC [1], but not after central axon injury [2]. We demonstrated that PPARα activity downstream of fatty acid synthesis in SGC contributes to promote axon regeneration in adult peripheral nerves and that the FDA approved PPARα agonist fenofibrate increased axon regeneration in the dorsal root, a model of poor sensory axon regeneration [1; 2]. Pathway analysis using KEGG demonstrated lipid metabolic pathways such as fatty acid and steroid metabolism along with PPAR signaling pathway in the top enriched pathways in all species (Fig. 5B-D). These observations further confirm the similarity between human and rodent SGC and suggest that PPARα activation is a promising therapeutic for nerve injuries and other pathological conditions of peripheral nerves.

### Human SGC express a greater variety of ion channels and receptors compared to rodent SGC

Astrocytes influence neural activity, in part by controlling the neuronal microenvironment through maintaining homeostasis of neurotransmitters, potassium buffering, and synaptic transmission. SGC express potassium channels and glutamate transporters, suggesting that they perform similar functions in the PNS. However, the exact composition of ion channels and receptors in human SGC is not well characterized. Pathway analysis of the SGC gene set in human SGC demonstrated an enrichment in ion channel function, specifically ion transport activity (Fig. 5D). We observed differences at the level of the classical SGC marker *Kcnj10* in mouse, which shows much lower expression in human, where we note substantially higher relative expression of *Kcnj3* (Fig 3D). Within the top enriched genes is SGC with channels and receptors functions, we detected 34 unique ion channel and receptors expressed in human SGC, with 14 of them in the top expressed genes in SGC (>100,000 normalized total counts, Table 2), 31 genes in mouse and 17 genes in rat (Fig. 6A, Supplementary Table 6). A majority of genes were related to potassium channels and glutamate receptors in both human and rodents (Fig. 6B). All species shared 5 genes; *Kcnj10, Gja1, Grid2* (Glutamate Ionotropic Receptor Delta Type Subunit 2), *Cacng4* (Calcium Voltage-Gated Channel Auxiliary Subunit Gamma 4) and *Aqp4* (Aquaporin-4) (Fig. 6A). While the potassium channel *Kcnj10* and the Gap junction *Gja1* have been previously reported as rodent SGC marker genes, *Grid2, Cacng4* and *Aqp4* were not. Interestingly, the glutamate receptor *Grid2* and the calcium voltage-gated channel *Cacng4* were enriched in fetal human astrocytes but not in mature astrocytes (Supplementary Table 2) [84]. Aquaporin-4 is a water channel predominantly found in astrocytes in the central nervous system and is believed to play a critical role in the formation and maintenance of the blood-brain barrier and in water secretion from the brain [51], further highlighting that the similarity between SGC and astrocytes is conserved in human.

**Table 2.** Top normalized counts expression of ion channels and receptors genes in human SGC

| Gene symbol | Channel type | Gene symbol | Receptor type |
| --- | --- | --- | --- |
| KCNE4 | Potassium channels | GRIK2 | Glutamate receptors |
| KCNH8 | Potassium channels | GRIN2B | Glutamate receptors |
| KCNJ3 | Potassium channels | CHRM3 | Cholinergic Receptors |
| KCNMB4 | Potassium channels | P2RY14 | Purinergic receptors |
| SCN7A | Sodium channels |  |  |
| SCN9A | Sodium channels |  |  |
| CACNA1D | Calcium channels |  |  |
| TRPM3 | TRP channels |  |  |
| ANO1 | Chloride channels |  |  |
| ITPR2 | Other |  |  |

**Figure 6:**
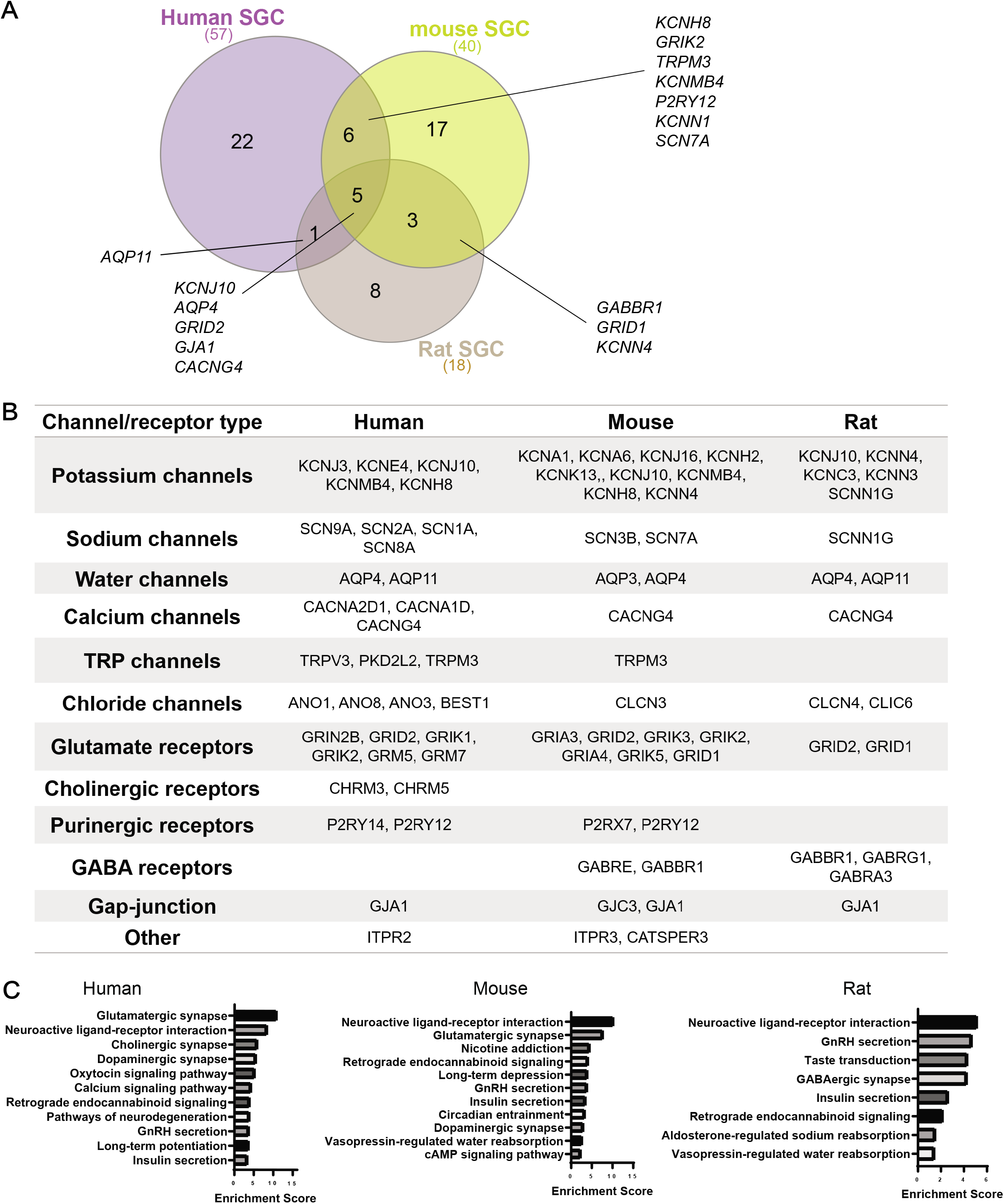
Enrichment of ion channels and receptors in human, mouse and rat SGC. (A) Venn diagram for enriched ion channel and receptors genes in SGC from mouse (40 genes), rat (18 genes) and human (57 genes). (B) Enriched ion channels and receptors in SGC compared to other cell types in the DRG (fold change threshold >1.5). (C) Pathway analysis (KEGG 2019) of enriched ion channels and receptors expressed in mouse, rat and human SGC.

Examination of enriched molecular pathways (KEGG) of the enriched channels and receptors gene sets in SGC for each species revealed that the top pathways were related to neuroactive ligand-receptor interaction, GnRH secretion, insulin secretion and endocannabinoid signaling pathway (Fig. 6C). Mouse and human SGC were also enriched for glutamatergic and dopaminergic synapse (Fig. 6C). The similarity in ion channel and receptor composition between human and rodent SGC suggests conserved physiological roles of SGC across species. However, the differences in subset of channels and receptors may inform future study design for therapeutic development targeting SGC in pain conditions and other peripheral neuropathies.

### Human SGC express SARS-CoV-2 coronavirus-associated factors and receptors

SGC have been implicated in pain conditions related to viral infection such as herpesvirus, varicella zoster virus and also swine hemagglutinating encephalomyelitis virus, which is related to the coronavirus family [25]. Current models suggest that SGC surrounding virally infected neurons may restrict the virus spread [25]. A recent study suggested that sensory neurons could be potential targets for the infection of SARS-CoV-2, with SARS-CoV-2 gaining access to the nervous system through entry into nociceptor nerve endings in the skin and luminal organs [67]. COVID-19, the disease caused by the SARS-CoV-2 can trigger many unexplained neurological effects including chronic pain. Price and colleagues found that a subset of human DRG neurons express the SARS-CoV-2 receptor angiotensin-converting enzyme 2 (ACE2) at the RNA and protein level [67]. DRG neurons also express SARS-CoV-2 coronavirus-associated factors and receptors (SCARFs), which were shown to be expressed in DRG at the lumbar and thoracic level as assessed by bulk RNA sequencing of human DRG tissue [67]. Having the resolution of single cell in human DRG, we assessed the expression of *Ace2* and *SCARF* genes specifically in SGC. While the *Ace2* receptor was lowly expressed in SGC in all three species, the assembly/trafficking factors *Rab10* and *Rab1a, Rab14, RhoA* and the restriction factors *Ifitim3* and *Ifitim2* were highly expressed (Fig. 7A,B, Table 3). Further examination of these abundant genes in human SGC revealed high similarities in expression across individual donors (Fig. 7B). Interferon-induced transmembrane proteins (IFITMs) restrict infections by many viruses, but a subset of IFITMs can enhance infections by specific coronaviruses. Recently, it has been showed that human IFITM3 with mutations in its endocytic motif enhances SARS-CoV-2 Spike-mediated cell-to-cell fusion and thus raised the concept that polymorphisms in IFITM3 can positively or negatively influence COVID-19 severity [66]. Weather IFITM3 expression in SGC enhance or limit viral infection in sensory ganglia remains to be determined. Together, these results suggest that SARS-CoV-2 may gain access to the nervous system through entry into sensory neurons at free nerve endings in organs and that SGC may attempt to restrict the local diffusion of the virus [25; 40].

**Figure 7:**
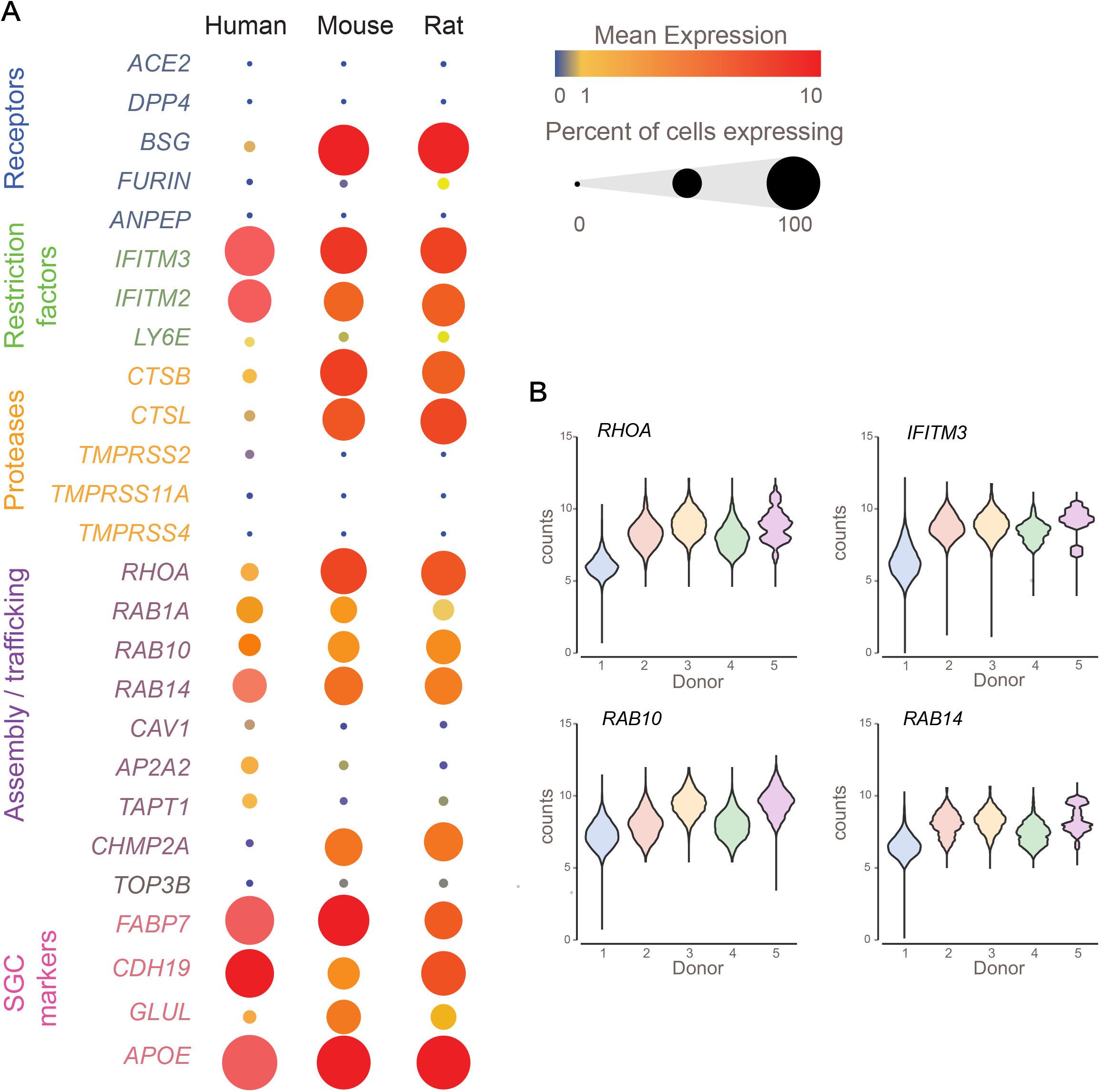
Enrichment of SCARF genes in human, mouse and rat SGC. (A) Bubble plot showing the normalized counts expression of SCARF genes in each SGC species. Dot size represent the percentage of cells expressing the corresponding gene and the color represents the mean expression level. (B) Violin plots of top normalized counts expression of SCARF genes in human SGC across individual donors.

**Table 3.** SCARF genes expression in human SGC

| **Gene symbol** | **Total counts** | **Gene product role** |
| --- | --- | --- |
| ACE2 | 727.00 | Receptor |
| DPP4 | 727.00 | Receptor |
| BSG | 21,943.62 | Receptor |
| FURIN | 3,753.20 | Proteases |
| ANPEP | 2,692.34 | Receptor |
| CD209 | 1,033.39 | Receptor |
| IFITM3 | 104,924.33 | Restriction factor |
| IFITM2 | 37,969.32 | Restriction factor |
| LY6E | 5,162.42 | Restriction factor |
| CTSB | 37,109.31 | Protease |
| CTSL | 18,138.03 | Protease |
| TMPRSS2 | 13,491.42 | Protease |
| TMPRSS11A | 4,456.63 | Protease |
| FURIN | 3,753.20 | Protease |
| TMPRSS4 | 1,013.52 | Protease |
| RHOA | 50,894.46 | Assembly / trafficking |
| RAB10 | 176,394.75 | Assembly / trafficking |
| RAB1A | 118,462.82 | Assembly / trafficking |
| RAB14 | 37,654.99 | Assembly / trafficking |
| CAV1 | 18,848.95 | Assembly / trafficking |
| AP2A2 | 50,665.45 | Assembly / trafficking |
| TAPT1 | 34,529.00 | Assembly / trafficking |
| CHMP2A | 8,086.96 | Assembly / trafficking |
| TOP3B | 5,275.81 | replication |
| FABP7 | 178,367.71 | SGC marker |
| CDH19 | 1,420,697.86 | SGC marker |
| GLUL | 56,518.68 | SGC marker |
| APOE | 550,114.38 | SGC marker |

## 4. DISCUSSION

The biology of SGC has remained poorly characterized under normal or pathological condition. Most of the current knowledge on SGC function stems from studies in rodents. Here we present a direct comparison of transcriptional profile of SGC in mouse, rat and human at the single cell level. Our findings suggest that key features of SGC in rodent models, such as similarities with astrocytes and enrichment for lipid metabolism and PPARα signaling, are conserved in human. However, notable differences exist in ion channels and receptors expression, which may suggest differences in SGC-neuron communication and functions in pain conditions and other peripheral neuropathies. Our study provides the potential to leverage rodent and human SGC properties and unravel novel mechanisms and potential targets for treating nerve injuries and other pathological conditions. Furthermore, depending on the specific target that is under study for therapeutic development, these parallels and differences should be considered in study design.

Our previous study demonstrated that SGC contribute to the nerve repair process in mice [1] and that the FDA approved PPARα agonist fenofibrate, which is used in dyslipidemia treatment, can increase axon regeneration after the dorsal root injury, a model of poor sensory axon growth [2]. Fenofibrate was surprisingly shown in clinical trials to have neuroprotective effects in diabetic retinopathy [5; 48] and traumatic brain injury [8]. Fenofibrate was also shown to exert analgesic and neuroprotective effects in rodent models of chronic neuropathic pain and inflammation as well as in some human studies [15; 53; 55]. The neuroprotective role of fenofibrate was also recently observed in a paclitaxel chemotherapy-induced peripheral neuropathy [7]. It is possible that this neuroprotection is mediated by an effect of PPARα activation in SGC. Together, these studies further support a central role for SGC in multiple pathological conditions affecting peripheral nerves. The observation that PPAR signaling is similarly enriched in human and rodents SGC opens the potential pharmacological repurposing of fenofibrate and that manipulation of SGC could lead to avenues to promote functional recovery after mechanical or chemical nervous system injuries.

Our study also highlights that the functional similarity of SGC and astrocytes is conserved across species. Both cell types undergo major changes under pathological conditions, which can have a protective function, but can also contribute to disease, and chronic pain [26]. One of the main functional similarities is buffering of extracellular potassium. In the CNS, glial buffering of extracellular potassium is carried out by astrocytes and consists of potassium uptake by inwardly rectifying potassium (Kir) channels [35]. Kir channels are key regulators of glial functions, which in turn determine neuronal excitability and axonal conduction [6]. Functionally, Kir channels can be divided into different subtypes based on their biophysical properties. Kir4.1 is an ATP-dependent potassium channel, whereas Kir3.1 is a G protein-activated potassium channel [6]. Most astrocytes express Kir4.1 but rat astrocytes and guinea-pig Muller glia have been shown to express Kir3.1 [54; 60]. Our studies revealed that human SGC preferentially express Kir3.1, mice SGC mainly express Kir4.1 and rat SGC express low level of both Kir channels and more SK3. Nerve damage was shown to downregulate the expression of Kir4.1 [72; 73; 79] and silencing Kir4.1 in the rat trigeminal ganglia leads to pain-like behavior [79]. Similarly, gain or loss of Kir4.1 affects astrocyte ability to regulate neuronal activity [11]. The diversity in potassium channel expression in SGC might suggest that differences in signaling mechanisms related to ATP or G protein coupled receptors define the physiological characteristics of SGC across species.

We also observed an enrichment for other types of channels in human SGC, including sodium and water channels, chloride channels and TRP channels. Transient receptor potential (TRP) proteins consist of a superfamily of cation channels that have been involved in diverse physiological processes in the brain as well as in the pathogenesis of neurological disease. TRP channels are widely expressed in the brain, including neurons and glial cells. Channels of TRP family have been shown to be involved in sensation and modulation of pain in peripheral ganglia but their expression in SGC have not been demonstrated. TRP channels contribute to the transition of inflammation and immune responses from a defensive early response to a chronic and pathological conditions. The expression of purinergic and glutamate receptor was also highly conserved between human and mice, suggesting that the ATP and glutamate dependent communication between neuron and glia in response to neuronal activity and pathological conditions is largely conserved. Whether different channels are expressed in SGC surrounding different types of sensory neurons remains to be determined. In mice we detected at least 4 SGC subtype with no evidence that one of the subtypes is dedicate to one class of sensory neurons [2]. Spatial transcriptomics approaches or in situ hybridization to molecularly characterize transcriptomes of DRG and their adjacent SGC [74] might shed light on the molecular properties of individual neuron-SGC units in human.

Another main feature common to astrocytes and SGC and conserved across species is the enrichment for genes related to lipid metabolism and expression of ApoE. In astrocyte, lipid metabolism is critical for synapse development and function *in vivo* [3; 76]. ApoE is predominantly secreted by astrocytes in the brain and functions as a major transporter of lipoproteins between cells. Of the three ApoE alleles, the ApoE4 allele is associated with an increased risk for Alzheimer’s disease (AD) [18; 42]. ApoE likely regulates AD risk in large part via effects on amyloid pathology [38]. However, several studies revealed a role for ApoE in lipid delivery for axon growth [39; 49; 77; 78]. We previously found that ApoE expression is increased in SGC via activation of the PPARα signaling after nerve injury [1]. The pathophysiological changes in AD are believed to arise in part from defects in neuronal communication in the central nervous system [32; 65]. However, decline in different sensory modalities are suggested to be a primary first-tier pathology [12]. In cultured sensory neurons, exogenously applied ApoE4 directly inhibits neurite outgrowth, whereas ApoE3 stimulates neurite outgrowth [50]. These studies suggest that the ApoE4 risk factors in human SGC may directly impact sensory neurons and potentially hearing dysfunction [27; 41; 43; 61; 64; 70; 75] and postural instability [4; 37], which have been associated with neurodegenerative disorders such as Alzheimer’s disease (AD) and age-related dementia in humans.

Our analysis of human SGC complements prior studies on human sensory ganglia that have characterized DRG samples in bulk sequencing or focused on expression of specific markers in sensory neurons [13; 52; 63; 74], revealing similarities and difference between mouse and human sensory neurons. Our study highlights the potential to leverage on rodent SGC properties and generate new knowledge that can be used to develop novel therapeutics to treat pain and other aspects of peripheral neuropathies.

## ACKNOWLEDGEMENTS

This research was funded by in part by a post-doctoral fellowship from The McDonnell Center for Cellular and Molecular Neurobiology to O.A, by NIH grant NS042595 to R.G, by The McDonnell Center for Cellular and Molecular Neurobiology to V.C., by a Pilot Project Award from the Hope Center for Neurological Disorders at Washington University to V.C. and by NIH grants NS111719, NS122260 and NS115492 to V.C. We gratefully acknowledge Greg Strout, Ross Kossina and Dr. James Fitzpatrick from the Washington University Center for Cellular Imaging (WUCCI), which is supported in part by Washington University School of Medicine, The Children’s Discovery Institute of Washington University, and St. Louis Children’s Hospital (CDI-CORE-2015-505 and CDI-CORE-2019-813) and the Foundation for Barnes-Jewish Hospital (3770) for assistance in acquiring and interpreting Transmission Electron Microscopy (TEM) data. The authors wish to thank the human tissue donor families for their generous donations to science, which made the human tissue work presented here possible. We thank Mid-America Transplant for providing access to donor tissue and their facilities, and J. Lemen for his time and surgical skill in assisting with hDRG extractions.

## CONFLICTS OF INTEREST

The authors declare no conflict of interest

## DATA AVAILABILITY

The raw Fastq files and the processed filtered count matrix for scRNA sequencing were deposited at the NCBI GEO database under the accession number GSE158892 (mouse), GSE169301 (rat and human). Data analysis and processing was performed using commercial code from Partek Flow package at https://www.partek.com/partek-flow/.

## AUTHOR CONTRIBUTIONS

O.A and V.C designed research and wrote the manuscript. O.A. performed mouse and rat single cell sequencing, bioinformatic analyses, immunofluorescence experiments and analyzed data. A.C. and L.Y. performed human single nucleus sequencing. R.F performed bioinformatic analyses and analyzed data. A.H collected rat DRG for single cell sequencing. A.M.M, R.W.G and V.C. supervised the project. All authors edited and approved the manuscript.

## SUPPLEMENTRAY FIGURES

**Supplementary Figure 1: Cell populations marker genes in human, mouse and rat DRG cells**

(A-C) t-SNE overlay for expression of cell population marker genes in human (A) mouse (B) and rat (C).

(D) t-SNE plots of combined 9,509 mouse cells from different data sets (Avraham et al. 2020 and Avraham et al, 2021) separated by color, unbiased clustering (Graph-based) and cell classification.

**Supplementary Figure: 2 Similarity of human SGC to Astrocytes and Schwann cells**

(A) VENN diagram for genes enriched in human SGC, mature and fetal Astrocytes

(B) Enriched human SGC genes compared to human mySC and nmSC

(C) Enriched human mySC and nmSC compared to mouse mySC and nmSC

(D) Enriched human SGC compared to subclusters of mouse SGC

## Notes

### Competing Interest Statement

The authors have declared no competing interest.

