## Supplementary figures and images for "Profiling the molecular signature of Satellite Glial Cells at the single cell level reveals high similarities between rodent and human"

### Supplementary Figure 1

#### Supplementary Figure 1

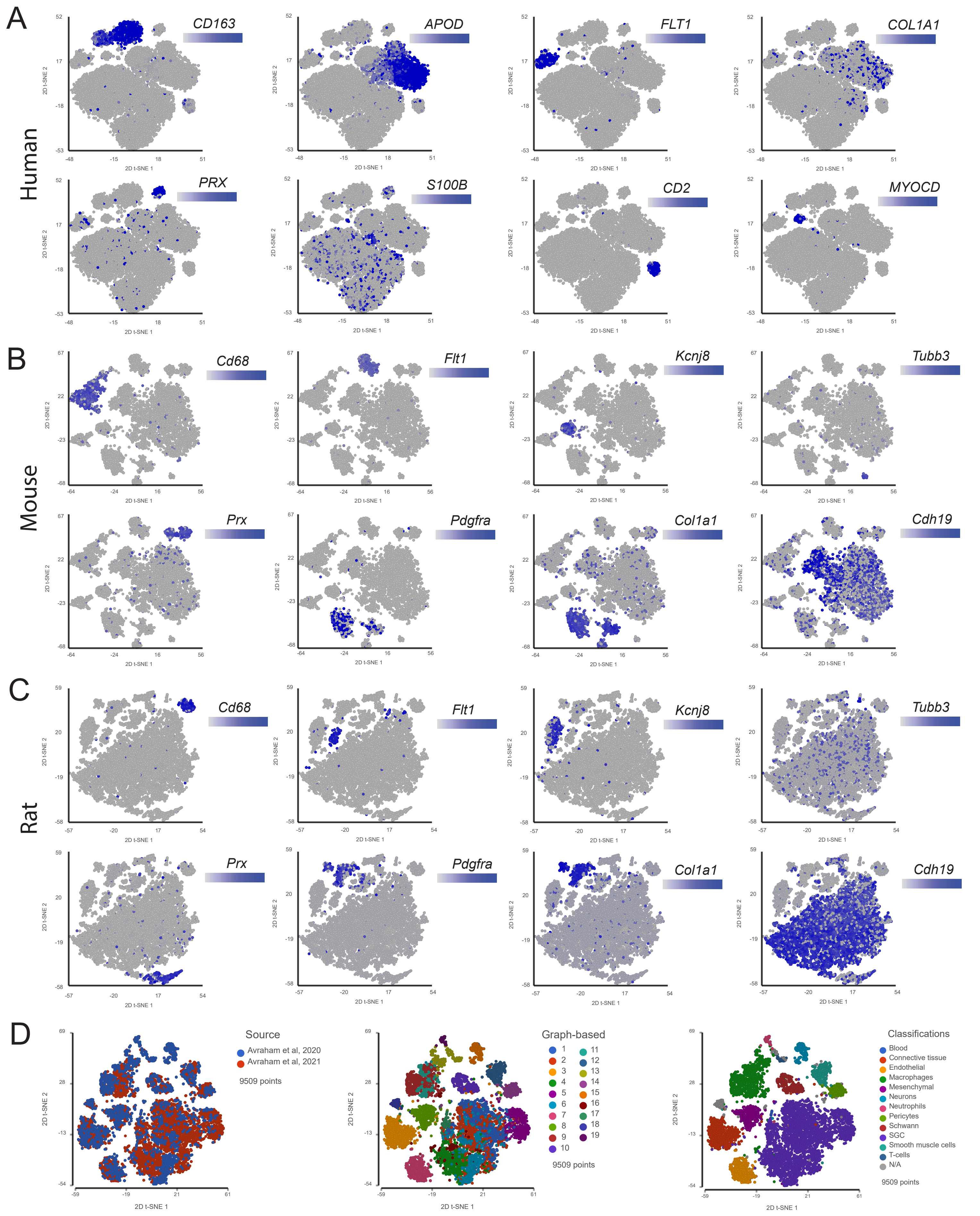

### Supplementary Figure 2

#### Supplementary Figure 2

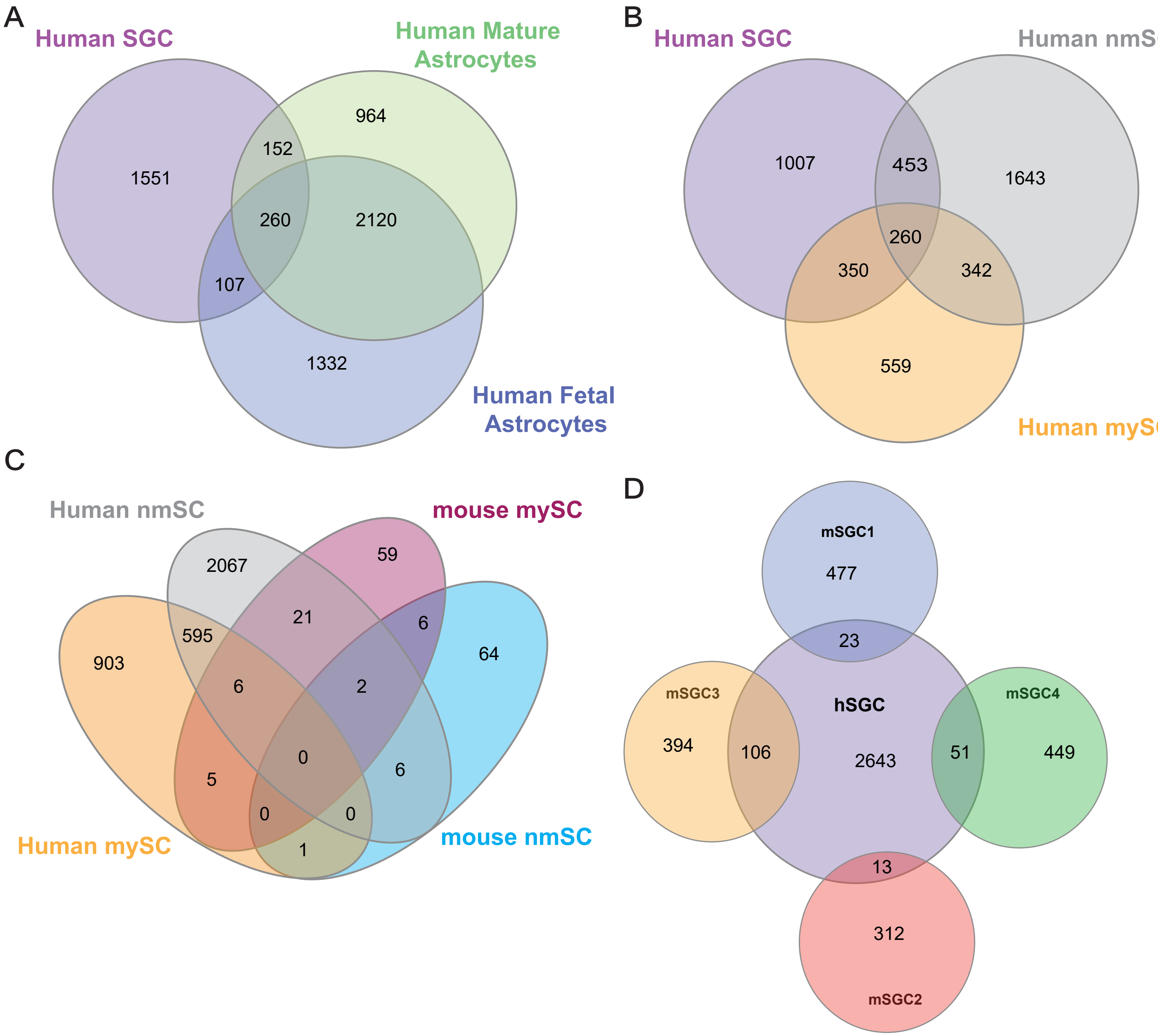
